# Endpoint and Epitope-specific Antibody Responses as Correlates of Vaccine-mediated Protection of Mice against Ricin Toxin

**DOI:** 10.1101/2020.05.06.081174

**Authors:** Greta Van Slyke, Dylan J Ehrbar, Jennifer Doering, Jennifer L. Yates, Ellen S.Vitetta, Oreola Donini, Nicholas J Mantis

**Affiliations:** Division of Infectious Disease, Wadsworth Center, New York State Department of Health, Albany, NY 12208; Department of Immunology and Microbiology, University of Texas Southwestern Medical School, Dallas, TX; Soligenix, Inc., Princeton, NJ, 08540

**Author notes:** Corresponding Author: Division of Infectious Disease, Wadsworth Center, 120 New Scotland Avenue, Albany, NY 12208. These authors contributed equally to this work (Greta Van Slyke, Dylan J Ehrbar).

**Keywords:** ricin, mouse, antibody, neutralizing, epitope, vaccine, biodefense

## Abstract

The successful licensure of vaccines for biodefense is contingent upon the availability of well-established correlates of protection (CoP) in at least two animal species that can then be applied to humans, without the need to assess efficacy in the clinic. In this report we describe a multivariate model that combines pre-challenge serum antibody endpoint titers (EPT) and values derived from an epitope profiling immune-competition capture (EPICC) assay as a predictor in mice of vaccine-mediated immunity against ricin toxin (RT), a Category B biothreat. EPICC is a modified competition ELISA in which serum samples from vaccinated mice were assessed for their ability to inhibit the capture of soluble, biotinylated (b)-RT by a panel of immobilized monoclonal antibodies (mAbs) directed against four immunodominant toxin-neutralizing regions on the enzymatic A chain (RTA) of RT. In a test cohort of mice (n=40) vaccinated with suboptimal doses of the RTA subunit vaccine, RiVax^®^, we identified two mAbs, PB10 and SyH7, which had EPICC inhibition values in pre-challenge serum samples that correlated with survival following a challenge with 10 x LD_50_ of RT administered by intraperitoneal (IP) injection. Analysis of a larger cohort of mice (n=645) revealed that a multivariate model combining endpoint titers and an epitope-profiling immune-competition capture (EPICC) assay values for PB10 and SyH7 as predictive variables had significantly higher statistical power than any one of the independent variables alone. Establishing the correlates of vaccine-mediated protection in mice represents an important steppingstone in the development of RiVax^®^ as a medical countermeasure under the United States Food and Drug Administration’s “Two Animal Rule.”

## 1. Introduction

The development of vaccines to counteract biothreats, including toxins, remains a high priority in many countries, including the United States [1, 2]. There are, however, formidable challenges to development and licensure [3]. Foremost is the need to assess vaccine efficacy (VE) in humans in the absence of clinical outcomes, because conventional efficacy studies are not ethical and field trials are not feasible for most if not all the Category A and B Biothreats. Under the Two Animal Rule, however, the United States Food and Drug Administration (FDA) will evaluate VE based on “adequate and well-controlled studies in animal models of the human disease or condition of interest” [4]. Of utmost importance are well-established and robust correlates of protection (CoP) that apply across species and and can be applied to humans. The undertaking of cross-species bridging studies has been successfully applied to anthrax vaccine adsorbed (AVA) to estimate survival probabilities in vaccinated human populations [5].

RT is classified by the Centers for Disease Control and Prevention (CDC) as a Category B biothreat, due to its extreme toxicity and ease by which it can be procured from castor beans (*Ricinus communis*), which are cultivated globally for industrial and cosmetic applications. In its mature form, RT is a ∼65 kDa heterodimeric glycoprotein consisting of an RNA N-glycosidase (RTA) joined by a disulfide to galactose/N-acetyl galactosamine (Gal/GalNAc)-specific lectin (RTB). RTB facilitates ricin endocytosis and retrograde transport from the plasma membrane to the endoplasmic reticulum (ER). Within the ER, RTA is released from RTB and retro-translocated into the cytoplasm, where it functions as a ribosome-inactivating protein (RIP) by depurinating a single residue in the sarcin-ricin loop (SRL) of 28S rRNA [6]. On the cellular level, ribosome arrest triggers the ribotoxic-stress response (RSR) and activates stress-activated protein kinase (SAPK) and programmed cell death (PCD) pathways [7]. The lethality of RT in animals translates into a multifaceted pathophysiology initiated by cellular damage and driven by inflammatory cytokines and death.

Historically, there are two advanced RTA-based subunit vaccines in development as a medical countermeasures (MCM) for RT intoxication, RV*Ec* and RiVax^®^ [8-10]. RV*Ec* is a truncated version of RTA that lacks the molecule’s C-terminal folding domain (residues 199-267), as well as a small hydrophobic loop in the N-terminus (residues 34-43) [11-13]. RV*Ec* was under development by the Department of Defense [8]. RiVax^®^ and is a full-length variant (267 residues) of RTA with point mutations at position Y80 to disrupt RTA’s RNA N-glycosidase activity and V76 that eliminates RTA’s ability to induce vascular leak syndrome (VLS) [14-16]. Pilot phase I clinical trials have indicated that both RV*Ec*- and RiVax^®^-adsorbed to aluminum salt adjuvants are safe and immunogenic in healthy adults [8, 10]. Moreover, the efficacy of the two vaccines has been demonstrated in animal models, including mice, and non-human primates (NHPs) [16-19]. In the case of RiVax^®^, for example, Rhesus macaques vaccinated three times by the intramuscular (IM) route at monthly intervals were immune to 3 x LD_50_ or aerosolized RT [18]. In mice, RiVax^®^ vaccination by intramuscular (IM), intraperitoneal (IP), subcutaneous (SC) and intradermal (ID) protects against hyper-lethal doses of RT administered by inhalation, gavage, or injection [19-22].

Despite the demonstrated pre-clinical efficacy of RiVax^®^ and RV*Ec*, a CoP for RT has not been formally established. Toxin-neutralizing antibody (TNA) titers are an obvious metric and are used as the universal standard in assessing immunity to tetanus and diphtheria toxins [23, 24]. Unfortunately, in the case of RT, TNA assays are insensitive, difficult to standardize, and of limited relevance to the primary cell types affected by RT *in vivo* (*e*.*g*., alveolar macrophages, Kupffer cells) [19]. However, it was reported that in a cohort of ∼300 mice vaccinated with RiVax^®^ by the intradermal (ID) or intramuscular (IM) routes that animals that survived lethal dose RT challenge had significantly higher RTA-specific serum IgG titers than those that died [25]. Other reports have noted a similar association [19, 22] but none have rigorously evaluated whether pre-challenge endpoint titers (EPT) are in fact predictive of survival.

In a recent study we raised the possibility that toxin-neutralizing, epitope-specific serum IgG titers might serve as a relative correlate of protection [22]. In the case of RT there is considerable evidence that toxin-neutralizing antibodies constitute only a fraction of the total antibody pool elicited following RiVax^®^ vaccination and that neutralizing antibodies target a limited number of immunodominant epitopes on RTA, referred to as epitope clusters I-IV [26]. In the past several years, we have amassed a collection of mAbs and camelid-derived single chain antibodies (V_H_Hs) against RTA that we have started to use as tools to investigate the polyclonal response elicited by RiVax^®^ antisera [18, 22, 26, 27]. Although we do not yet have a full understanding of the relative importance of epitope-specific antibodies in contributing to vaccine-induced immunity, passive protection studies in mice and NHP have indicated that single mAbs or combinations of mAbs are sufficient to afford immunity to levels similar to those achieved through active vaccination [28-31].

In this report, we explored EPT as well as values derived from an epitope profiling immune-competition capture (EPICC) assay as possible correlates of vaccine-mediated protection of mice against a lethal RT challenge dose by IP injection. The results demonstrate that EPT and EPICC values each afforded significant power in predicting survival, but that a multivariate model combining both metrics (i.e., EPT and EPICC values) improved predictability. The EPICC assay has the benefit to being species neutral and therefore potentially applicable across species as a possible co-correlate of immunity to RT.

## 2. Material and Methods

### 2.1 Chemicals and biological reagents

RT; *Ricinus communis* agglutinin II; RCA_60_) and biotin (b)-RT were purchased from Vector Laboratories (Burlingame, CA). A thermostable batch of RiVax^®^ was provided by Soligenix, Inc (Princeton, NJ), as described [18]. The murine mAbs used in this study were affinity purified from serum-free hybridoma supernatants using protein G chromatography at the Dana Farber Cancer Institute Monoclonal Antibody Core facility (Boston, MA). Unless noted otherwise, chemicals were obtained from the Sigma-Aldrich Company (St. Louis, MO).

### 2.2 Mouse vaccination and RT challenge studies

Mouse experiments were conducted in strict accordance with protocol #18-384 approved by the Wadsworth Center’s Institutional Animal Care and Use Committee (IACUC). The Wadsworth Center is an American Association for Laboratory Animal Science (AALAS) accredited institution. Female BALB/c mice (IMSR Cat# TAC:balb, RRID:IMSR_TAC:balb; 8-12-week old) were purchased from Taconic Biosciences (Rensselaer, NY) and housed at the Wadsworth Center under conventional specific pathogen-free conditions. Mouse vaccinations and RT challenges were carried out as reported previously [22]. Mice were immunized with RiVax^®^ (0.3, 1 or 3 µg) administered SC on days 0 (prime) and 21 (boost) [22]. Sera were harvested by submandibular bleeding on day 30. Mice were challenged by IP injection with 5 x LD_50_ RT (∼1 *μ*g/mouse; 10 *μ*g/kg) on study day 35 and monitored for 7 days thereafter for morbidity and weight loss.

### 2.3 ELISA

Indirect and competitive capture ELISAs were performed as described [22]. Nunc Maxisorb F96 microtiter plates (Thermo Scientific, Pittsburgh, PA) were coated with RT (1 *μ*g/ml in PBS, pH 7.4) overnight at 4°C. The plates were blocked with phosphate buffered saline (PBS) containing goat serum (2% v/v; Gibco, Grand Island, NY) and Tween-20 (0.1% v/v). Serum samples were serially (1:2) diluted in blocking solution. Horseradish peroxidase (HRP)-labeled goat anti-mouse IgG polyclonal antibodies (SouthernBiotech, Birmingham, AL) were used as secondary antibodies. TMB (3,3′,5,5′-tetramethylbenzidine; Kirkegaard & Perry Labs, Gaithersburg, MD) was used as colorimetric detection substrate and reactions were stopped with 1 N phosphoric acid. Plates were read on a SpectroMax 250 spectrophotometer and analyzed with Softmax Pro 5.4.5 software (Molecular Devices, Sunnyvale, CA). End-point titers were defined as the dilution where absorbance >3-times above the background *(e.g*., blank wells). Seroconversion was defined as an endpoint titer in a RT ELISA of ≥ 1:50. Geometric mean titers (GMTs) were calculated from the endpoint titers. Mice that had not seroconverted, as determined by ELISA, were assigned a GMT of 1 for the purposes of statistical analysis.

### 2.3 Epitope profiling immune-competition capture (EPICC) assay

Immulon 4HBX 96-well microtiter plates (Thermo Scientific) were coated with indicated anti-RTA mAbs (1 µg/ml in 0.1 mL) in PBS, pH 7.4 at room temperature for 1 h and then blocked overnight at 4°C, as noted above. For EPICC, the amount of b-RT used in the assay was adjusted to the EC_90_ for each MAbs (30-200 ng/ml) [32], as shown in **Table 1**. Serial dilutions of control or immune sera were mixed with b-RT (EC_90_) in duplicate in PVC microtiter plates, incubated for 15 min and then transferred using a multichannel pipette to Immulon 4HBX 96-well microtiter plates (Thermo Scientific) coated with indicated anti-RTA mAbs (1 µg/ml in 0.1 mL). in PBS, pH 7.4 at room temperature for 1 h and then blocked overnight at 4°C, as noted above. As controls, each mAb (10 µg/ml) was competed with itself to establish a 100% inhibitory baseline. The microtiter plates were incubated at RT for 1 h, washed and then probed with streptavidin-HRP (1ug/ml; Thermo Scientific) and developed with 3,3′,5,5′-Tetramethylbenzidine (TMB) (Kirkegaard & Perry Labs). Plates were analyzed with a SpectroMax 250 spectrophotometer using Softmax Pro 5.2 software (Molecular Devices).

**Table 1.**
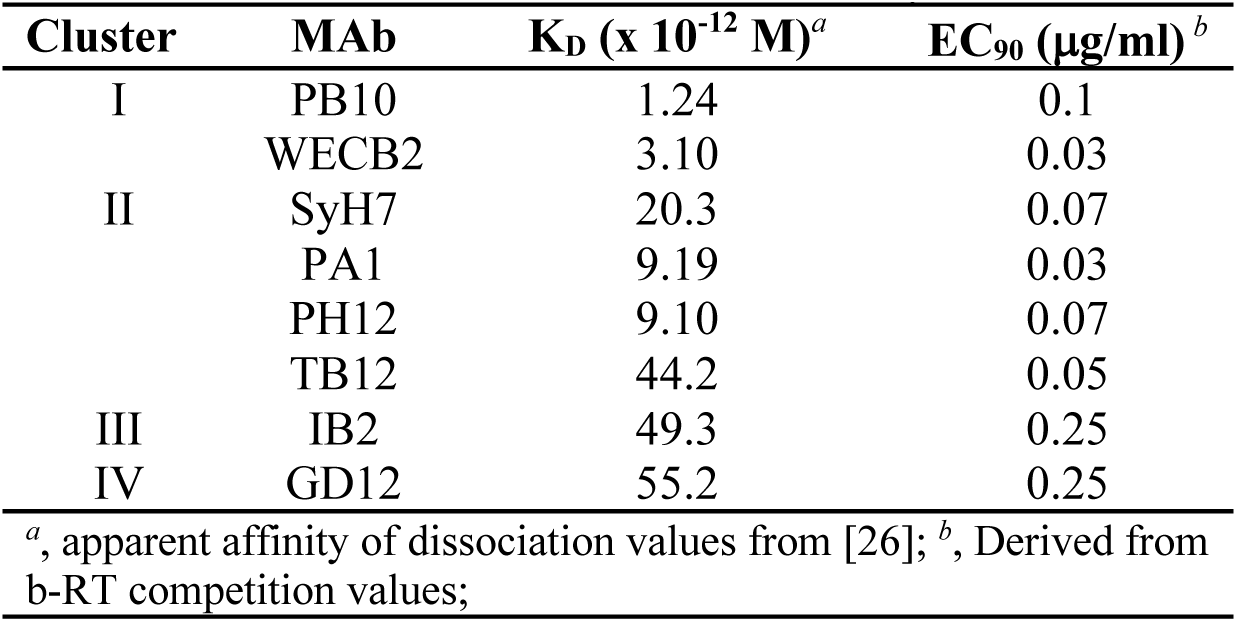
Mouse mAbs used for EPICC analysis.

Inhibition of RT binding was calculated as a percentage of b-RT binding to the coated MAb, where: [100-(OD_450_*C*/OD_450_*B*)*100]= % RT binding inhibited by competitor, and *C*= competed well, *B*= b-R (EC_90_). Pooled sera from RiVax-vaccinated rabbits and Rhesus macaques [18] were diluted in PBS prior to use for EPICC analysis.

### 2.3 Statistical analysis

Endpoint titers were log-transformed prior to statistical analysis. Endpoint titers were compared using one-way analysis of variance (ANOVA) followed by Dunnett’s multiple comparison test. Survival data were tested using the log rank Mantel-Cox test. In all cases the significance threshold was set at P < 0.05. ANOVA and log rank tests were performed using GraphPad Prism v. 8.0 for Windows (GraphPad Software, San Diego, CA, USA).

Correlations between endpoint titers or EPICC values and mouse survival following challenge with RT were determined by simple logistic regression. In the initial cohort of 40 mice, the optimal set of predictive variables was defined using least absolute shrinkage and selection operator (LASSO) penalized logistic regression [33]. Ten-fold cross-validation was performed 10 times to select the optimal values for the *λ* penalty parameter. All sets of values for each separate potential correlate of protection were standardized to a mean of 0 and a standard deviation of 1 before LASSO regression. The selected variables were then measured in the mice included in subsequent experiments and included in analysis of the larger dataset of 645 mice.

The predictive performance of each model constructed from the larger dataset was assessed by receiver operating characteristic (ROC) analysis, which allowed us to examine the trade-off between sensitivity and specificity along the entire range of values for each variable. The area under the curve (AUC) was also used as a measure of the predictive value of each variable, while Delong’s test was used to compare the AUC of each model [34]. P-values resulting from Delong’s test were adjusted for multiple comparisons with the Holm-Sidak method. Analyses were performed in R 3.4.2 [35], the R package pROC for ROC analysis [36], and the R package glmnet for LASSO penalized logistic regression [37].

## 3. Results

### 3.1 Probing RiVax^®^ with mAbs directed against four spatially distinct toxin-neutralizing epitope clusters

We previously described four spatially distinct immunodominant B cell epitope clusters (I-IV) on RiVax^®^ [26, 27, 38]. Each cluster is defined by one or more RT-neutralizing mAbs (**Table 1**). Cluster I, defined by PB10, is focused around RTA’s *α*-helix B (residues 94-107), a protruding element previously known to be a target of potent RT-neutralizing antibodies [39, 40]. Cluster II, defined by SyH7 and PA1, is located on the back side of RTA, relative to the active site pocket [26]. Cluster III is targeted by MAb IB2 and is in close proximity to RTA’s active site [41]. Finally, Cluster IV, defined by GD12, forms a diagonal sash from the front to back of the subunit. We previously reported that antisera from NHPs and humans vaccinated with RiVax^®^ competes with mAbs from cluster I (PB10) and cluster II (SyH7, PA1) for binding to RT [18].

We performed cross-competition capture ELISAs with each of the eight representative mAbs as confirmation that the four epitope clusters are spatially distinct (**Table 1**). In this modified capture ELISA, microtiter plates were coated with individual mAbs and probed with soluble b-RT, in the absence or presence of a competitor mAb. A reduction in the capture of soluble b-RT in the presence of a competitor was interpreted as epitope overlap or steric hindrance [42]. We refer to this modified competition ELISA as EPICC.

As shown in **Figure 1**, there was across the board intra-cluster competition, with only limited inter-cluster competition. For example, cluster II mAbs, SyH7, PA1, PH12 and TB12, competed with each other, but not with mAbs in clusters I, III, or IV. Similarly, IB2 (cluster III) competed with itself but not with the other seven mAbs. The exception was competition between GD12 (cluster IV), and two cluster I mAbs, PB10 and R70. This cross-cluster competition (I versus III) is attributed to the contact of GD12 with the *α*-helix B [26]. JD4, the other cluster IV mAb in our collection, did not compete with PB10 or R70. These results demonstrate our ability to “interrogate” specific immunodominant epitope clusters on RiVax^®^ by competition ELISA.

**Figure 1.**
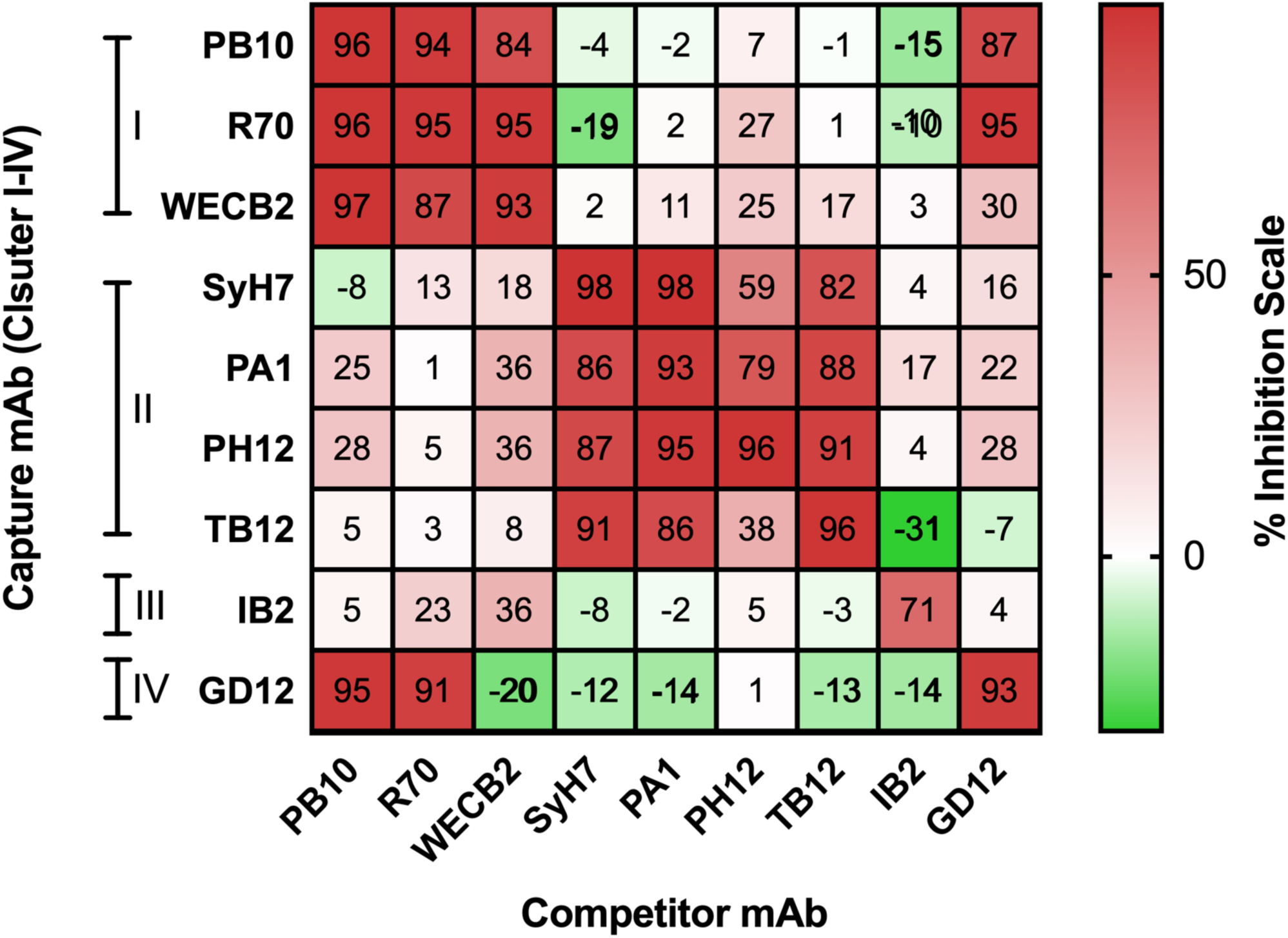
Spatial segregation of immunodominant neutralizing B cell epitopes on RTA, as determined by EPICC. Cross competition ELISAs with indicated mAbs used as capture (y-axis) or competitor (x-axis) in solution with b-RT. The heatmap scheme indicates percent inhibition of competitor MAb, as compared to b-RT alone with inset numbers indicating percent reduction in a representative experiment. Negative values indicate enhancement of b-RT-in capture.

To examine whether the epitope-specific antibodies against the 4 clusters identified in mice are also present in other species, we performed EPICC analysis with pooled immune sera from rabbits and Rhesus macaques that had been vaccinated with RiVax™. As shown in **Figure 2**, pooled sera from all three species competed with the murine mAbs representing clusters I (PB10, WECB2), II (PA1, SyH7, PH12, TB12), III (IB2), and IV (GD12). These results demonstrate that the four immunodominant epitope clusters on RTA identified in mice are also targets of antibodies in other species.

**Figure 2.**
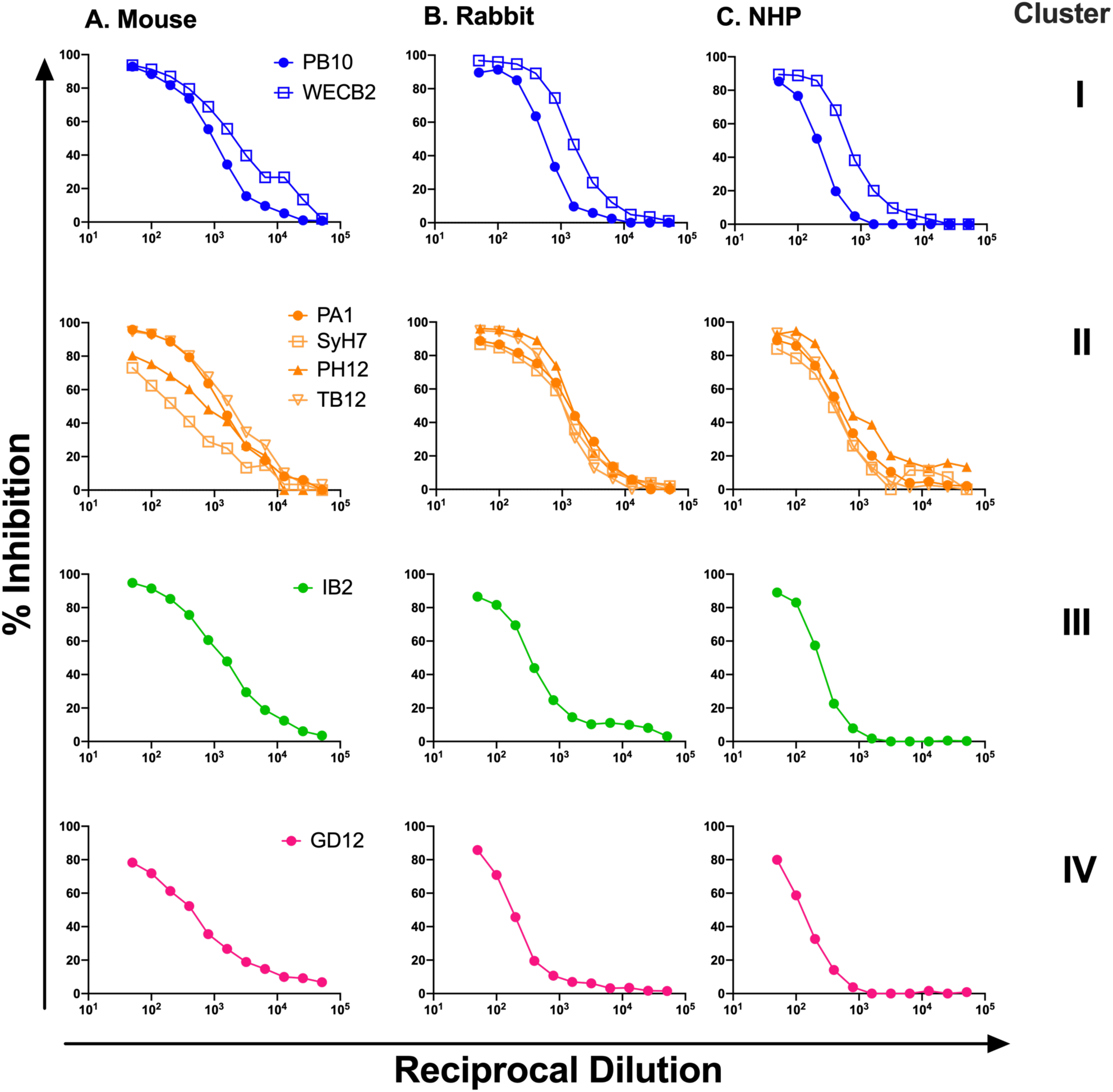
EPICC analysis of antisera from rabbits and NHPs vaccinated with RiVax. Pooled polyclonal sera from RiVax^®^ -vaccinated (A) mice, (B) rabbits and (C) Rhesus macaques were subjected to EPICC with mAbs (see inset legends in first column) directed against RTA epitope clusters I, II, III and IV.

### 3.2 Preliminary analysis of EPT and EPICC values as correlates of vaccine-mediated protection

We next investigated the relative importance of RT-specific serum IgG titers and EPICC values as predictors of survival following a lethal injection of RT. To generate a preliminary data set, a group of 40 mice were divided into two cohorts and vaccinated SC on days 0 and 21 with RiVax^®^ at optimal (1 µg; cohort 1) or sub-optimal (0.3 µg; cohort 2) doses. Serum samples were collected on day 30, and the mice were challenged on day 35 with 5 x LD_50_ of RT administered by IP injection. Mice were then monitored for 7 days post-challenge for weight loss and morbidity, as described [38]. The results of each mouse in that experiment are presented in **Appendix 1**.

In cohort 1, 19 of the 20 vaccinated mice (95%) survived RT challenge, whereas in cohort 2 only 9 out of 20 survived (45%). As shown in **Figure 3**, pre-challenge, RT-specific EPT were significantly higher in mice that survived RT exposure (n=12), as compared to the decedents (median = 4.408 log_10_ transformed EPT for the survivors vs. 3.204 for the decedents, *U* = 72.50, P = 0.0033, as determined by two-tailed Mann-Whitney *U* test). In terms of pre-challenge EPICC analysis, mice that survived RT challenge had significantly higher PB10 (median = 29.4% inhibition for survivors vs. 9.8% for decedents, *U* = 65, P = 0.0017), SyH7 (median = 10.2% for survivors vs. 4.95% for decedents, *U* = 97, P = 0.0362), and PH12 (median = 16.43% for survivors vs. 4.782 for decedents, *U* = 94, P = 0.0286) inhibition values as compared to the decedents. In contrast, EPICC values for IB2 (P = 0.1631) and GD12 (P = 0.6308) were not significantly different between groups of mice that survived or died (**Figure 3**).

**Figure 3.**
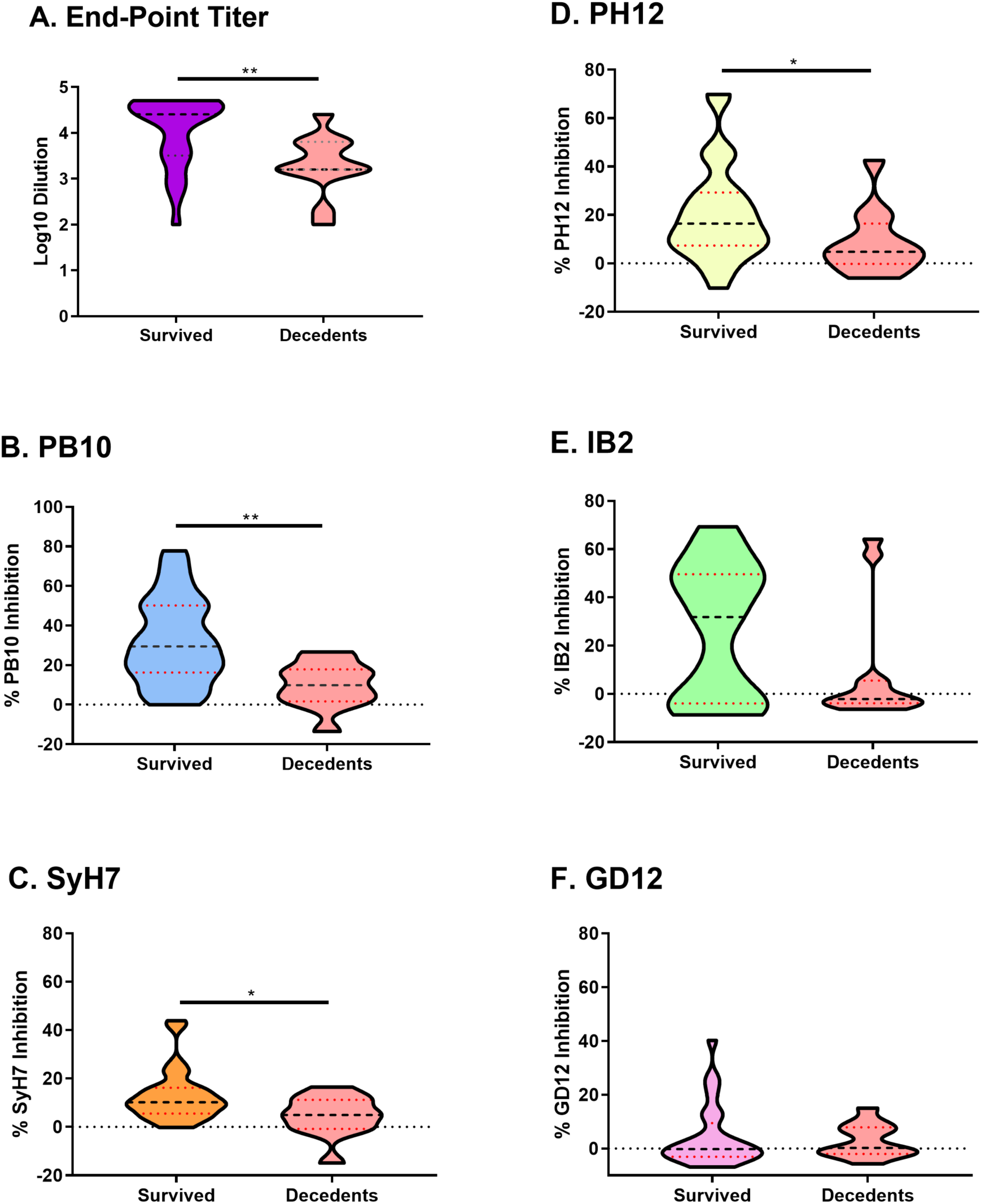
Pilot study reveals EPT and EPICC as putative correlates of protection in a cohort of RiVax^®^ vaccinated mice. Female BALB/c mice (n=40) were vaccinated with suboptimal or near optimal doses of RiVax^®^ (0.3 or 1 *μ*g) at days 0 and 21 and challenged with 10 x LD_50_ RT on day 35. Sera was collected from mice on day 30 and analyzed by (A) indirect ELISA to determine EPT and (B-F) EPICC analysis. Statistical significance between survivors and decedents is denoted by an asterisk (unpaired t test; p < 0.02).

**Figure 4.**
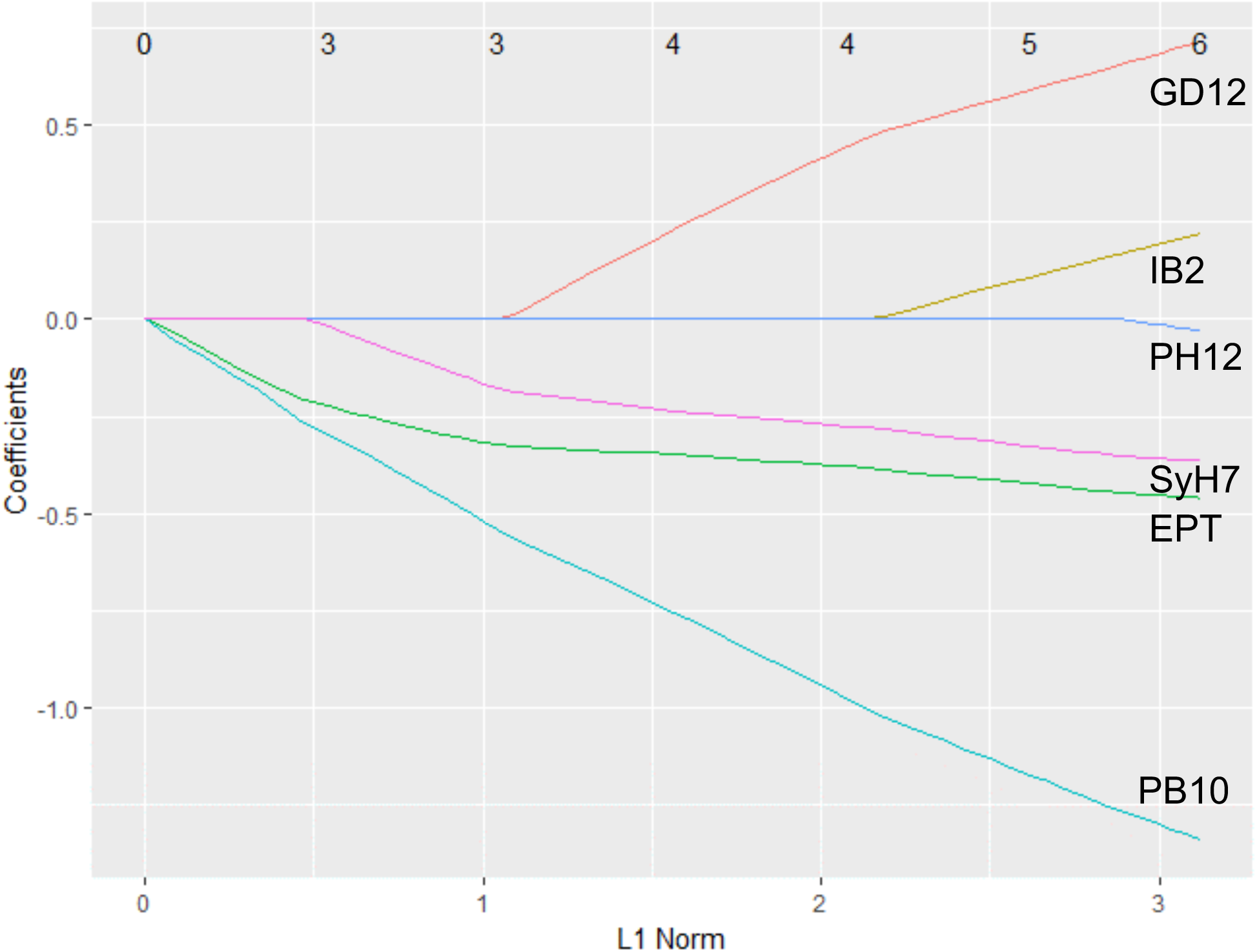
EPT and EPICC values for SyH7 and PB10 as correlates of immunity to RT. Results presented in Figure 3 were subjected to variable selection using LASSO penalized logistic regression. LASSO coefficient profiles were generated for all potential correlates of protection: EPT, PB10, SyH7, PH12, IB2 and GD12. Each curve corresponds to a potential predictive variable with coefficients plotted against the L1 Norm regularization term (lower x-axis). The upper x-axis depicts the number of nonzero coefficients at the respective regularization parameter.

We next generated a series of univariate logistic regression models to further examine the relationships between survival and pre-challenge, RT-specific serum EPT and EPICC. Logistic regression demonstrated a significant relationship between EPT and survival (pP < 0.01). For the EPICC analysis, inhibitory levels of each of the five RTA-specific mAbs was designated as the independent variable and death was designated as the dependent variable. For PB10 and SyH7, there was a significant correlation between EPICC values and survival, whereas for PH12, IB2, and GD12 there was not. To assess the predictive performance of the logistic regression models we employed ROC analysis. The models yielded AUC values of 0.7842 for EPT, 0.8065 for PB10 inhibition and 0.7113 for SyH7 inhibition. LASSO penalized logistic regression selected the optimal set of predictive variables as being EPT, combined with PB10 and SyH7 EPICC inhibition values. Thus, combining EPT and EPICC values derived from PB10 or SyH7 was tentatively the most effective predictor of immunity to a lethal dose of injected RT.

### 3.4 A multivariant model combining EPT and EPICC as a correlate of vaccine-mediated protection

We next examined a much larger cohort of mice (n=645) that had been uniformly vaccinated with RiVax^®^ on days 0 and 21, and then challenged with a 5x LD_50_ dose of RT on day 35 [22]. Serum samples were collected from the animals five days prior (day 30) to the challenge. Within this cohort, 374 mice survived ricin challenge and 271 died (**Appendix 2**).

In accordance with the preliminary study, RT-specific IgG titers in sera collected on day 30 were higher in the mice that subsequently survived a lethal dose, as compared to mice that succumbed (median = 3.607 log_10_ transformed EPT for survivors vs. 1.699 for decedents, *U* = 12245, P < 0.0001, as determined by two-tailed Mann-Whitney *U* test; **Figure 5A**). Similarly, EPICC revealed that mice that survived the challenge had significantly higher PB10 (median = 41.95% inhibition for survivors vs. 3.244 for decedents, *U* = 17659, P < 0.0001) and SyH7 inhibition values (median = 19.70% for survivors vs. 1.001% for decedents, *U* = 19069, P < 0.0001), as compared to the decedents (**Figure 5 B,C**). Moreover, all 3 variables correlated with survival when examined in the single-variable models (P < 2e-16 for all) (**Table 3**; **Figure 6**). The univariate EPT model had the highest AUC of the three (0.8792, 95% CI 0.8526-0.9057), followed by PB10 (0.8258, 95% CI 0.7939-0.8576) and SyH7 (0.8119, 95% CI 0.779-0.8447). The AUC values for PB10 and SylH3 were each were significantly lower than the AUC derived from EPT (Holm-Šídák adjusted P < 0.05, as determined by Delong’s test), demonstrating that when considering univariate analysis EPT is superior. However, a multivariate model considering EPT and EPICC for PB10 and SyH7 as predictive variables had a significantly higher AUC (0.901, 95% CI 0.8773-0.9246) than did EPT alone (Holm-Šídák adjusted P < 0.05) (**Table 3**). In summary, EPT and EPICC values for both PB10 and SyH7 have the potential to serve as co-correlates of vaccine-mediated protection of mice against RT with the term “co-correlate” being defined by Plotkin as “…a quantity of a specific immune response to a vaccine that is 1 of >2 correlates of protection and that may be synergistic with other correlates.” [23]

**Table 2.**
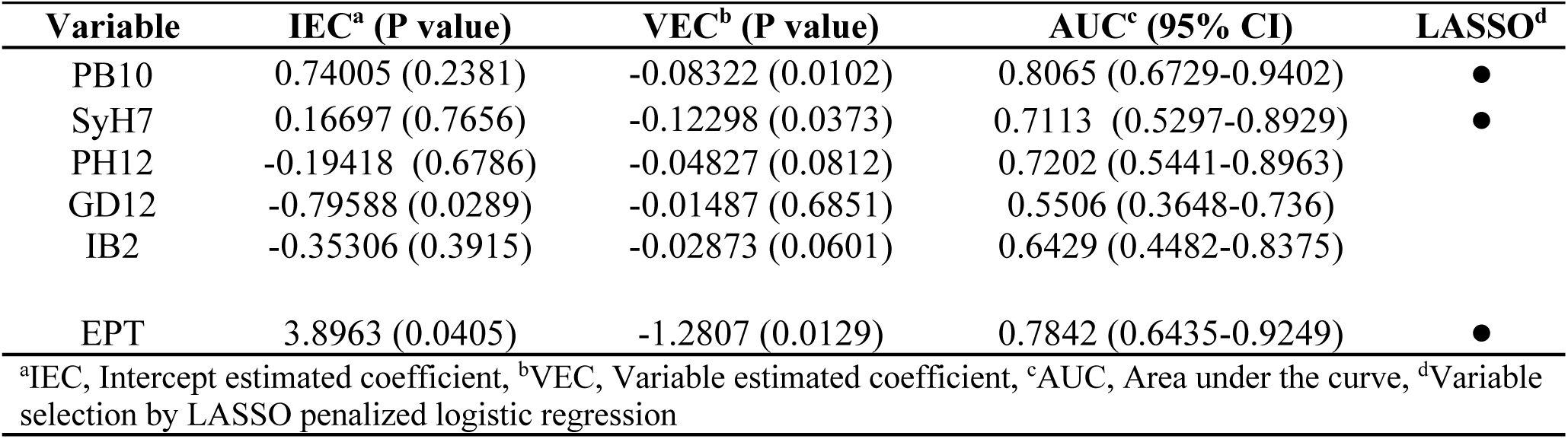
Logistic regression analysis of EPICC and endpoint titers.

**Table 3.**
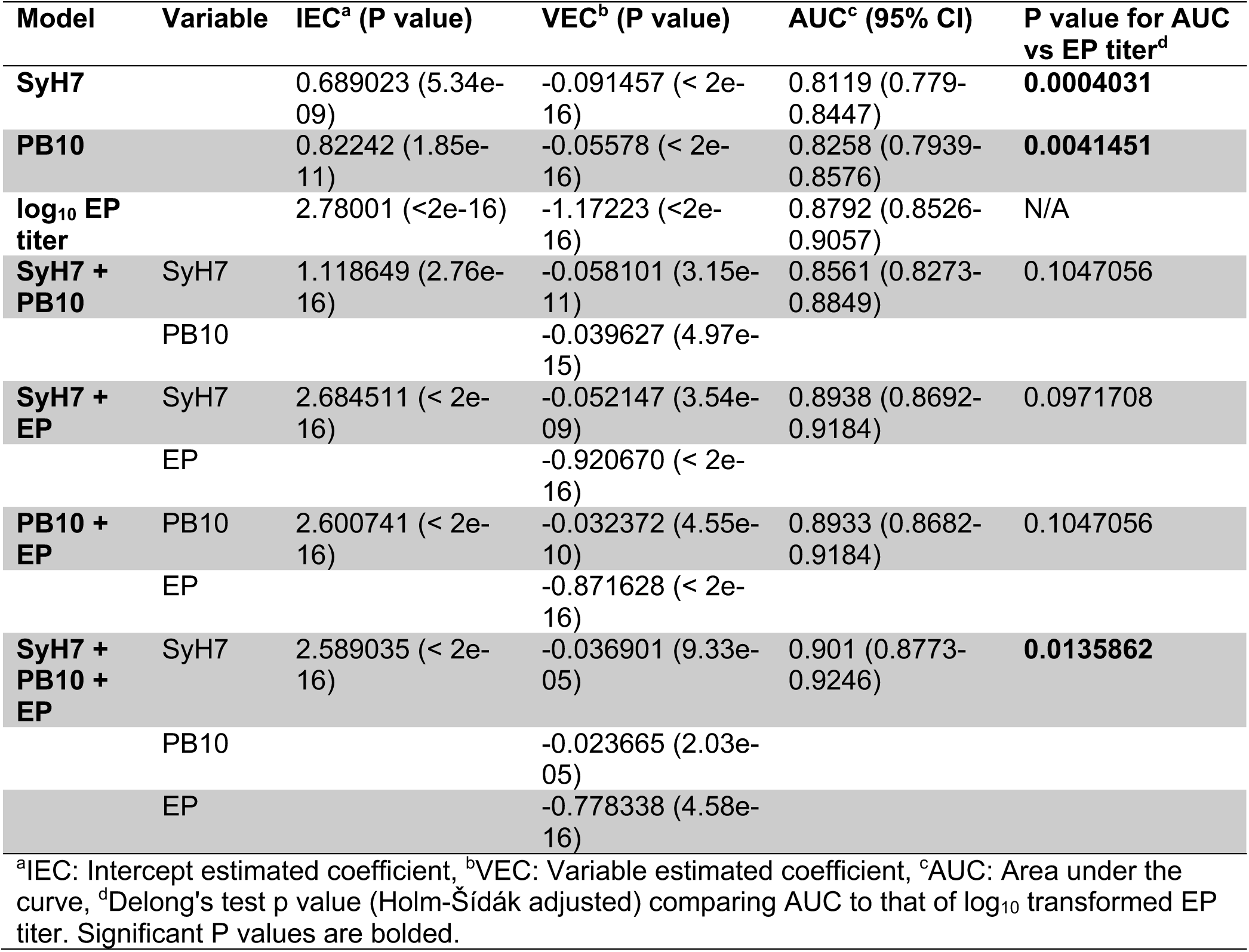
Model comparison as immune correlates of protection against RT in mice.

|  |  |  |  |  |  |
| --- | --- | --- | --- | --- | --- |
| Model | Variable | IEC<sup>a</sup> (P value) | VEC<sup>b</sup> (P value) | AUC<sup>c</sup> (95% CI) | P value for AUC vs EP titer<sup>d</sup> |
| **SyH7** |  | 0.689023 (5.34e-09) | -0.091457 (< 2e-16) | 0.8119 (0.779-0.8447) | **0.0004031** |
| **PB10** |  | 0.82242 (1.85e-11) | -0.05578 (< 2e-16) | 0.8258 (0.7939-0.8576) | **0.0041451** |
| **log<sub>10</sub> EP titer** |  | 2.78001 (<2e-16) | -1.17223 (<2e-16) | 0.8792 (0.8526-0.9057) | N/A |
| **SyH7 + PB10** | SyH7 | 1.118649 (2.76e-16) | -0.058101 (3.15e-11) | 0.8561 (0.8273-0.8849) | 0.1047056 |
|  | PB10 |  | -0.039627 (4.97e-15) |  |  |
| **SyH7 + EP** | SyH7 | 2.684511 (< 2e-16) | -0.052147 (3.54e-09) | 0.8938 (0.8692-0.9184) | 0.0971708 |
|  | EP |  | -0.920670 (< 2e-16) |  |  |
| **PB10 + EP** | PB10 | 2.600741 (< 2e-16) | -0.032372 (4.55e-10) | 0.8933 (0.8682-0.9184) | 0.1047056 |
|  | EP |  | -0.871628 (< 2e-16) |  |  |
| **SyH7 + PB10 + EP** | SyH7 | 2.589035 (< 2e-16) | -0.036901 (9.33e-05) | 0.901 (0.8773-0.9246) | **0.0135862** |
|  | PB10 |  | -0.023665 (2.03e-05) |  |  |
|  | EP |  | -0.778338 (4.58e-16) |  |  |
| <sup>a</sup>IEC: Intercept estimated coefficient, <sup>b</sup>VEC: Variable estimated coefficient, <sup>c</sup>AUC: Area under the curve, <sup>d</sup>DeLong's test p value (Holm-Šidák adjusted) comparing AUC to that of log<sub>10</sub> transformed EP titer. Significant P values are bolded. |  |  |  |  |  |

**Figure 5.**
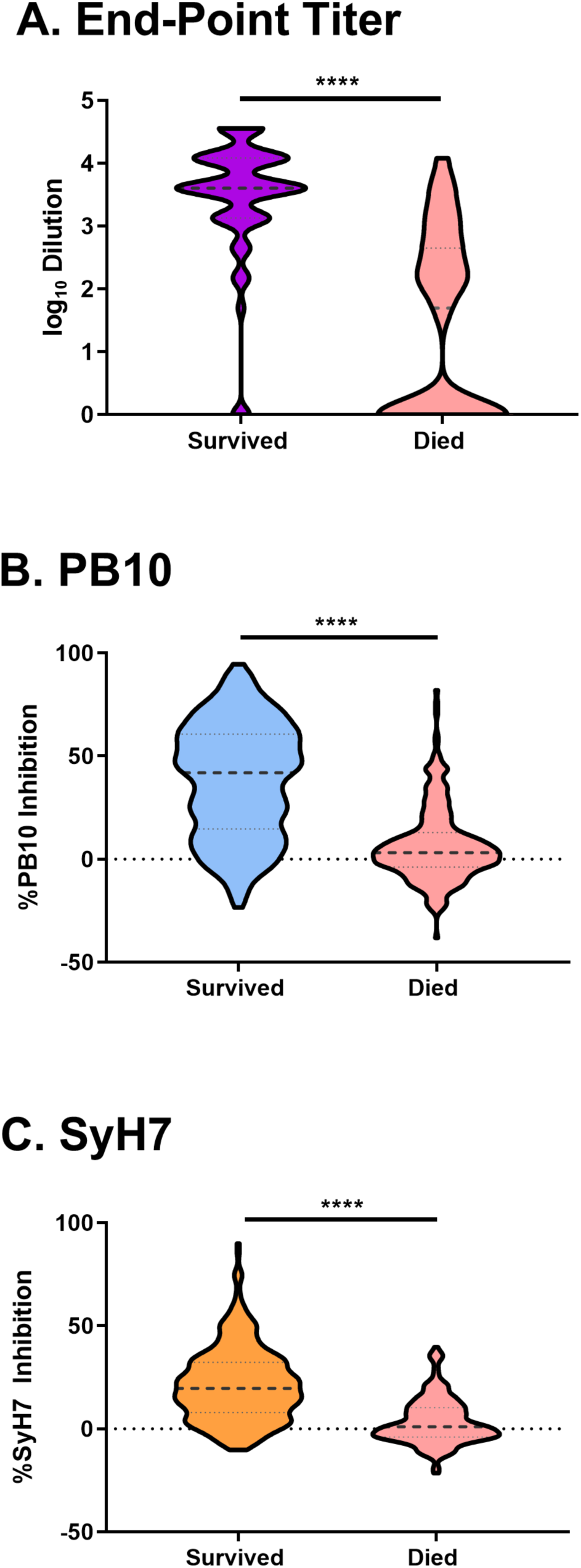
EPT and EPICC as putative correlates of protection in a large cohort of RiVax^®^ vaccinated mice. A cohort of 646 mice total vaccinated with RiVax^®^ (dose range 0.3-3ug) on days 0 and 21 were challenged with RT on day 35, as reported in **Appendix 2**. Violin plots of pre-challenge serum samples examined for (A) EPT and (B-C) EPICC values with PB10 and SyH7. Statistical significance between survivors and decedents is denoted by an asterisk (Mann-Whitney test; p>0.0001).

**Figure 6.**
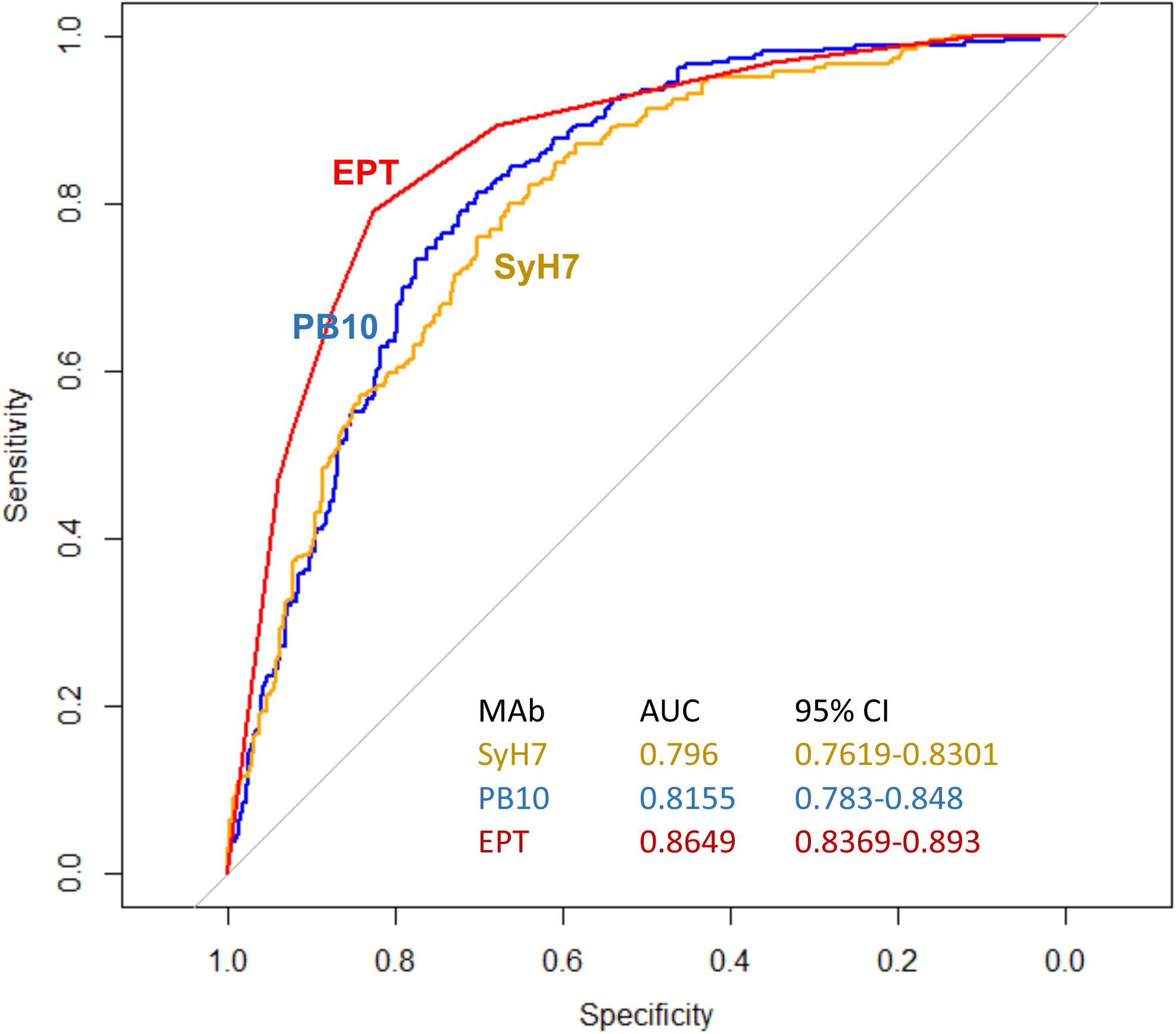
ROC curve analysis of predictive performance of classifying variables in the large cohort. Curves are based on univariate logistic regression models including the labeled variable. Area under the curve (AUC) values and corresponding 95% confidence intervals are included for each predictive variable.

## 4. Discussion

In this report we investigated in a mouse model the potential of pre-challenge, RT-specific serum EPTs, as well as RTA epitope-specific antibody levels, to serve as CoP from lethal dose ricin toxin challenge, administered by injection. In both a pilot and larger cohort of mice, EPT emerged as having significant predictive value as a CoP. This finding unto itself constitutes an advance in the development of RiVax^®^ considering that EPTs have never been formally been evaluated for its prognostic use with survival as an endpoint. By the same token, EPICC values derived using two mAbs, PB10 and SyH7, directed against different immunodominant toxin-neutralizing epitopes on RTA, were also significantly predictive of survival in the mouse model. Combining EPT and RTA-specific epitope reactivity, as determined by EPICC, ultimately provided the highest predictive power.

It should be noted that one advantage of the EPICC assay over direct competition ELISAs is that it is species neutral, because it relies on the detection of captured biotinylated ricin toxin using avidin-HRP, rather than using species-specific secondary reagents. This enables direct cross-species comparisons in pre-clinical testing. Another advantage is that EPICC involves the capture soluble antigen (RT), rather than detection of antigen immobilized on polystyrene microtiter wells. This is biologically significant because RT and its individual toxin subunits are partially denatured on plastic surfaces, resulting in the exposure of cryptic epitopes and possibly perturbation of native secondary structures.

We have recently initiated studies in Rhesus macaques to evaluate the bridging of potential of EPT and EPICCs as indicators of RiVax^®^ -induced protection against RT. As alluded to above, it was previously demonstrated that Rhesus macaques that received three IM vaccinations with RiVax^®^ (100 *μ*g/dose) on days 0, 30, and 60 were protected against a 3 x LD_50_ toxin challenge by aerosol on day 110 [18]. In that study, serum samples were assessed for RT-specific EPT, TNA, and preliminary competition ELISAs with a subset of RTA-specific mAbs, including PB10 and SyH7. However, establishing a CoP from that experiment alone was not possible considering that all animals survived ricin challenge (except for one that died from causes unrelated to toxin exposure). Nonetheless, that report and the work presented in F**igure 2** of this study bodes well for the applicability of EPICC to the NHP model, in that the four RT-neutralizing immunodominant epitopes on RTA identified in mice appear to be shared across species. Whether the specific EPICC profiles will translate across species is under evaluation. In our current study, PB10 and SyH7 inhibition values, representing epitope clusters I and II, elicited following RiVax vaccination were predictive of survival. However, in preliminary analysis of NHP samples, IB2, representing cluster III, and not PB10 or SyH7 appear to correlate with protection (G. Van Slyke, D Ehrbar, C. Roy E. Vitetta, N. Mantis, unpublished results).

It should be noted that the conclusions drawn from our current study are limited to a single vaccination route (SC) and a single challenge route (IP). It is possible that specific CoPs for RT may depend on the site of vaccination and the mode of toxin exposure. Inhalation of RT triggers a particularly complex pathophysiology that results in acute lung injury (ALI) and acute respiratory distress syndrome (ARDS) involving multiple different cell types and inflammatory cytokines [43-45]. It is unclear where and how antibodies function to protect the lung against RT-induced damage, although a recent report from our group demonstrated that a humanized version of PB10 is sufficient, when given prophylactically by intravenous infusion, to protect Rhesus macaques against aerosolized ricin [31]. Thus, there are at least some parallels between rodents and NHPs, considering that PB10 has comparable toxin-neutralizing efficacy in mice as NHPs [46].

In summary, we have described a multivariate model combining EPT and EPICC values that affords high confidence in predicting survival of mice following a lethal challenge with RT. This model should facilitate the development of RiVax^®^ as a MC for RT under the United States Food and Drug Administration’s “Animal Rule.”

## Abbreviations

ELISA: Enzyme-linked immunosorbent assay
EPICC: epitope profiling immune-competition capture
mAb: Monoclonal antibody
b: biotinylated
RT: ricin toxin
RTA: RT A chain

## Acknowledgements

The work described in this manuscript was supported by Contract No. HHSN272201400039C (to OD/RS; Soligenix, Inc.) and grant AI125190 (to NJM) from the National Institutes of Allergy and Infectious Diseases (NIAID), National Institutes of Health (NIH) and the Simmons-Patigian Chair (EV) at UT Southwestern. The content is solely the responsibility of the authors and does not necessarily represent the official views of the NIH. The funders had no role in study design, data collection and analysis, decision to publish, or preparation of the manuscript.

## Author Contributions (CRediT)

**GVS, NJM**: conceptualization; **GVS, DJE, JW, and JY**: investigation; **GVS, DJE, DJV, EV, OD, NJM**: formal analysis; **GVS, DJE NJM**: writing original draft: **EV, OD, NJM**: writing, review and editing; **NJM:** Supervision and project administration.

## Declaration of Interest statement

GVS, DJE, JW, JY, EV, and NJM declare no competing interests. OD is an employee of Soligenix, Inc. which holds the license for RiVax^®^.

